# Cardiolipin targets a dynamin related protein to the nuclear membrane

**DOI:** 10.1101/2020.10.01.322057

**Authors:** Usha Pallabi Kar, Himani Dey, Abdur Rahaman

## Abstract

Dynamins are large cytoplasmic GTPases that are targeted to specific cellular membranes which they remodel via membrane fusion or fission. Although the mechanism of target membrane selection by dynamins has been studied, the molecular basis of conferring specificity to bind specific lipids on the target membranes is not known in any of the family members. Here, we report a mechanism of nuclear membrane recruitment of Drp6 that is involved in nuclear remodeling in *Tetrahymena thermophila*. Recruitment of Drp6 depends on a domain that binds to cardiolipin-rich bilayers. Consistent with this, the nuclear localization of wildtype Drp6 was inhibited by depleting cardiolipin in the cell. Cardiolipin binding was blocked with a single amino acid substitution (I553M) in the membrane-binding domain of Drp6. Importantly, the I553M substitution was sufficient to block nuclear localization without affecting other properties of Drp6. Consistent with this result, co-expression of wildtype Drp6 was sufficient to rescue the localization defect of I553M variant in *Tetrahymena*. Inhibition of cardiolipin synthesis or perturbation in Drp6 recruitment to nuclear membrane caused defects in the formation of new macronuclei post-conjugation. Taken together, our results elucidate a molecular basis of target membrane selection by a nuclear dynamin, and establish the importance of a defined membrane-binding domain and its target lipid in facilitating nuclear expansion.

## INTRODUCTION

Topological changes and remodeling of membranes are fundamental processes in cellular physiology. Intricate biological machineries have evolved to facilitate these changes in living cells. Dynamins and dynamin related proteins (DRPs) comprise a family of large GTPases that catalyze membrane remodeling reactions (Praefcke and McMahon 2004). Members of the dynamin family are mechano-chemical enzymes which couple the free energy of GTP hydrolysis with membrane deformation, thereby performing important cellular functions ranging from scission of membrane vesicles, cytokinesis, and maintaining mitochondrial dynamics to conferring innate antiviral immunity (Ramachandran and Schmid 2018). The common features shared by all dynamins and DRPs are the presence of a large GTPase domain (GD), a middle domain (MD) and a GTPase effector domain (GED), which distinguish them from other GTPases (Praefcke and McMahon 2004). The MD and the GED are involved in oligomerization and regulation of GTPase activity (Ramachandran, Surka et al. 2007). The feature that distinguishes DRPs from classical dynamins is the lack of a defined pleckstrin homology domain (PH domain) and a proline rich domain (PRD).

All dynamin proteins undergo rounds of assembly and dis-assembly on the target membrane, and tubulate the underlying membrane, which is required for fission or fusion function. The members of this family become associated with lipids on the target membrane and are important for performing cellular functions (Ramachandran and Schmid 2018).The PH domain in classical dynamins binds to phosphatidyl inositol 4,5 bis-phosphate (PIP2) at the target sites of endocytosis, and plays an essential role in vesicle scission during endocytosis (Zheng, Cahill et al. 1996). All the DRPs lack PH domains and instead possess either lipid binding loops or trans-membrane domains for membrane recruitment or association (Ramachandran and Schmid 2018). A stretch of positively-charged amino acid residues in the lipid binding loops of all known DRPs interacts with the negatively-charged head groups of the lipids and this interaction is important for target membrane association (Rujiviphat, Meglei et al. 2009, von der Malsburg, Abutbul-Ionita et al. 2011, Bustillo-Zabalbeitia, Montessuit et al. 2014, Smaczynska-de, Marklew et al. 2019, Wang, Guo et al. 2019, Yan, Qi et al. 2020).

Nuclear remodeling including its expansion is a fundamental process in eukaryotes, the mechanism of which is not well understood in any organism. It requires remodeling of nuclear membrane and incorporation of new materials including lipids and proteins into the existing membrane. *Tetrahymena thermophila*, a unicellular ciliate, exhibits nuclear dimorphism. Each cell harbors a small, diploid, transcriptionally inactive micronucleus (MIC) and a large, polyploid, transcriptionally-active macronucleus (MAC) (Karrer 2000). During sexual conjugation in *Tetrahymena*, two cells of complementary mating types first form a pair, followed by a series of complex nuclear events resulting in the loss of the parental MAC. Subsequently, new MIC and MAC are formed through the fusion of haploid nuclei produced from parental micronuclei. The precursors of the new MIC and MAC are identical in size and in genome content at the initial stage of nuclear differentiation. However, two of the four new MICs rapidly enlarge, making the final volume 10-15 fold larger, and develop into MACs (Cole, Cassidy-Hanley et al. 1997). This process calls for a sudden and dramatic expansion of nuclear envelope. DRP6, which is one of the eight dynamin-related protein paralogs in *Tetrahymena*, is specifically upregulated when the MICs rapidly expand to form new MACs. Inhibition of Drp6 function results in a profound deficiency in the formation of new MACs (Rahaman, Elde et al. 2008). It has been recently demonstrated that Drp6 functions as an active GTPase and self-assembles into higher order helical spirals and ring structures, and therefore resembles other members of the family (Kar, Dey et al. 2018).

In the present study, we have elucidated a mechanism for DRP6 recruitment in the nuclear membrane. Our results reveal that Drp6 directly interacts with membrane lipids, and that cardiolipin acts as its physiologically-important target lipid. Further, we have identified a lipid-binding domain in Drp6, and provide evidence that the domain plays a pivotal role in nuclear association of Drp6 and therefore in nuclear expansion.

## RESULTS

### A DRP targeting domain (DTD) is important for nuclear recruitment of Drp6

All the known dynamin family proteins perform cellular functions by associating with target membrane. Classical dynamins contain a pleckstrin homology (PH) domain responsible for membrane binding (Fig. 1a). Drp6 associates with nuclear envelope and regulates nuclear remodeling in *Tetrahymena*. However, Drp6 like other DRPs lacks a PH domain or any recognizable membrane binding domain (Fig. 1a). Sequence alignment revealed the presence of a region in Drp6 that is located at the position of the PH domain of classical dynamin (Fig. 1b, S1A) and that was earlier named the Drp targeting determinant (DTD) (Elde, Morgan et al. 2005).

**Fig. 1.**
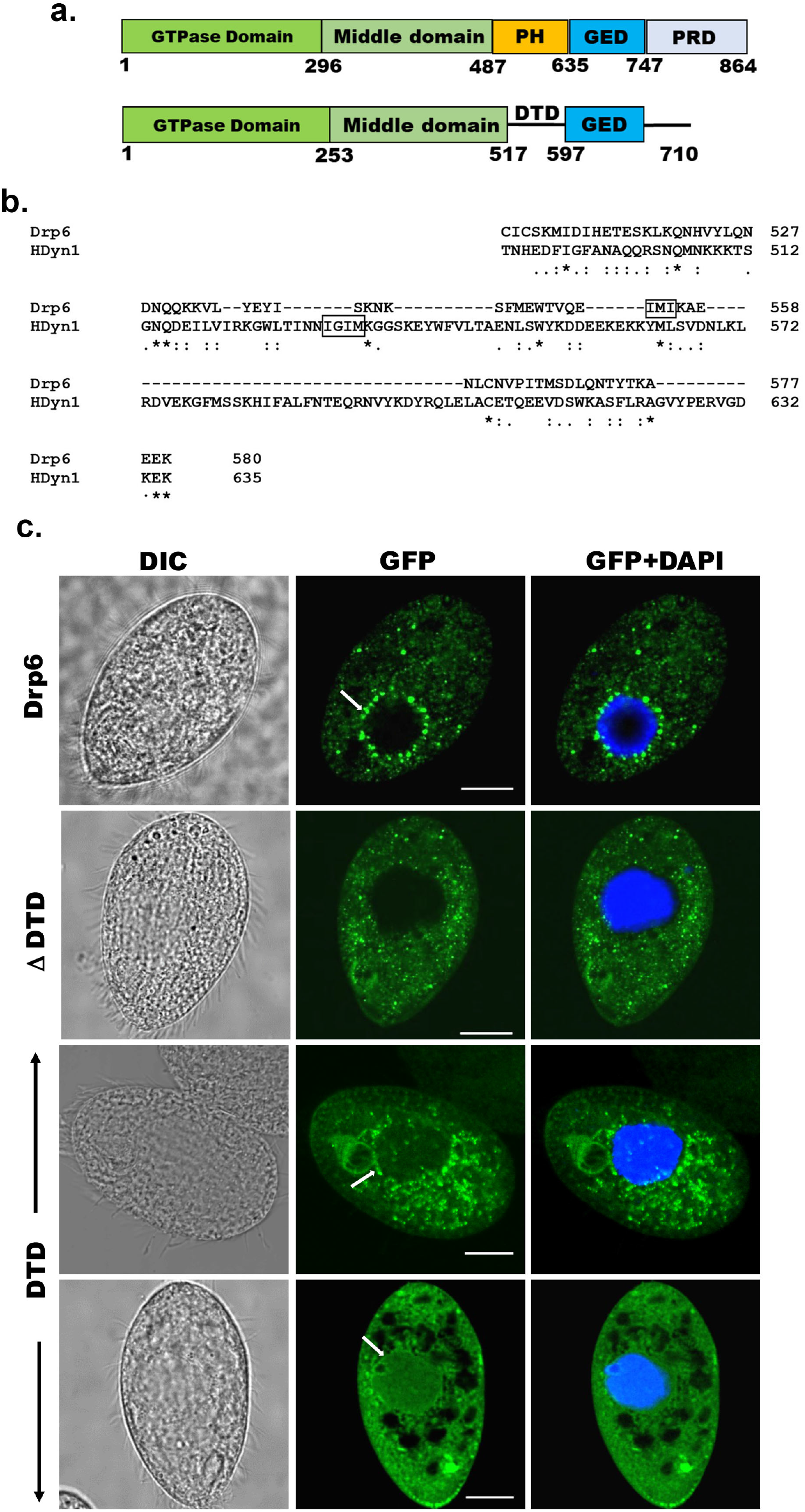
Identification of the region of Drp6 important for nuclear recruitment. a. Diagram showing domains of human dynamin 1 (Top) and Drp6 (Bottom). Five domains of dynamin indicated as G domain, Middle domain, PH domain, GED and PRD. Drp6 contains 3 domains but lack PH domain and PRD. Numbers indicate the position of amino acids in the protein. b. Sequence alignment of *Tetrahymena* Dynamin Related Protein 6 (Drp6) and human dynamin1 (HDyn1) generated using Clustal Omega. Only the PH domain of HDyn1 and the corresponding aligned region of Drp6 are shown. The hydrophobic patch (IGIM) of PH domain important for membrane insertion is shown within a box. A putative hydrophobic patch (IMI) in Drp6 is also within box. c. Confocal images of fixed *Tetrahymena* cells after DAPI staining. Cells expressing GFP-Drp6 (Drp6), GFP-Drp6ΔDTD (ΔDTD), and Drp6-DTD (DTD) are shown. DAPI in blue marks the nucleus. Localization on the nuclear envelope is indicated by arrow. Bar = 10μm.

To assess if the DTD is important for recruitment of Drp6 to target membranes, full length *DRP6* and *drp6*Δ*DTD* were expressed separately as N-terminal GFP fusion proteins and their cellular distributions were compared by confocal microscopy. As expected, Drp6 was chiefly present on the nuclear envelope and also on some cytoplasmic puncta (Fig. 1c and S1B). These cytoplasmic puncta of Drp6 are ER-derived vesicles (Rahaman, Elde et al. 2008). When confocal images of *GFP-drp6ΔDTD* expressing cells were analyzed, it was observed that the deletion of DTD resulted in complete loss of nuclear localization, and it was mainly associated with cytoplasmic puncta (Fig. 1c and S1B). These results clearly demonstrate that DTD is necessary for recruiting Drp6 to the nuclear envelope but not to cytoplasmic puncta. We next evaluated if the DTD is sufficient for nuclear envelope targeting, by expressing it as a GFP-fusion. The confocal images of cells expressing GFP-*drp6-DTD* showed that GFP-DTD often appeared as cytoplasmic puncta, but that nuclear envelope localization in a subset of cells is detectable albeit less prominently as compared to that of GFP-Drp6 (Fig. 1c and S1B). This suggests that the DTD is able to interact with the nuclear envelope. This interaction appears weaker than for the full-length protein, suggesting that other domains also contribute to nuclear recruitment of Drp6. Similarly, other dynamin family members rely for their targeting on a membrane-binding domain but also other domains such as GTPase domain (Vallis, Wigge et al. 1999, von der Malsburg, Abutbul-Ionita et al. 2011, Bramkamp 2012). We conclude that DTD is essential but not sufficient for Drp6 recruitment to the nuclear envelope. In contrast, Drp6 does not require its DTD for targeting to the ER-derived cytoplasmic vesicles

### DTD is the membrane-binding domain of Drp6

Recruitment of a protein to a target membrane is achieved either by interaction with membrane lipids or by forming a complex with another membrane protein. The DTD is essential for recruitment of Drp6 to the nuclear envelope, and may represent a membrane-binding domain. To test this idea, we generated N-terminal histidine tagged- *drp6-DTD* and *DRP6* for bacterial expression and purification, and then used the purified proteins in lipid overlay assays. Drp6 was purified to near homogeneity (Fig. 2a) and incubated with total *Tetrahymena* lipid spotted on nitrocellulose membrane either in the presence or absence of GTP. Drp6 interacts with *Tetrahymena* lipid with or without GTP (Fig. 2b) suggesting that it harbors a lipid-binding domain. To identify the lipids with which Drp6 interacts, we performed the overlay assay using commercially available strips spotted with fifteen different lipids (Echelon Biosciences, USA). The results demonstrated that Drp6 specifically interacts with three phospholipids, namely phosphatidylserine (PS), phosphatidic acid (PA) and cardiolipin (CL) (Fig. 2b). In order to find out whether lipid binding is a property of the DTD, we partially purified DTD as an N-terminal his tagged protein (Fig. 2a) and used it for the overlay assay. Like the full-length protein, DTD also interacts with all three phospholipids (Fig. 2b).

**Fig. 2.**
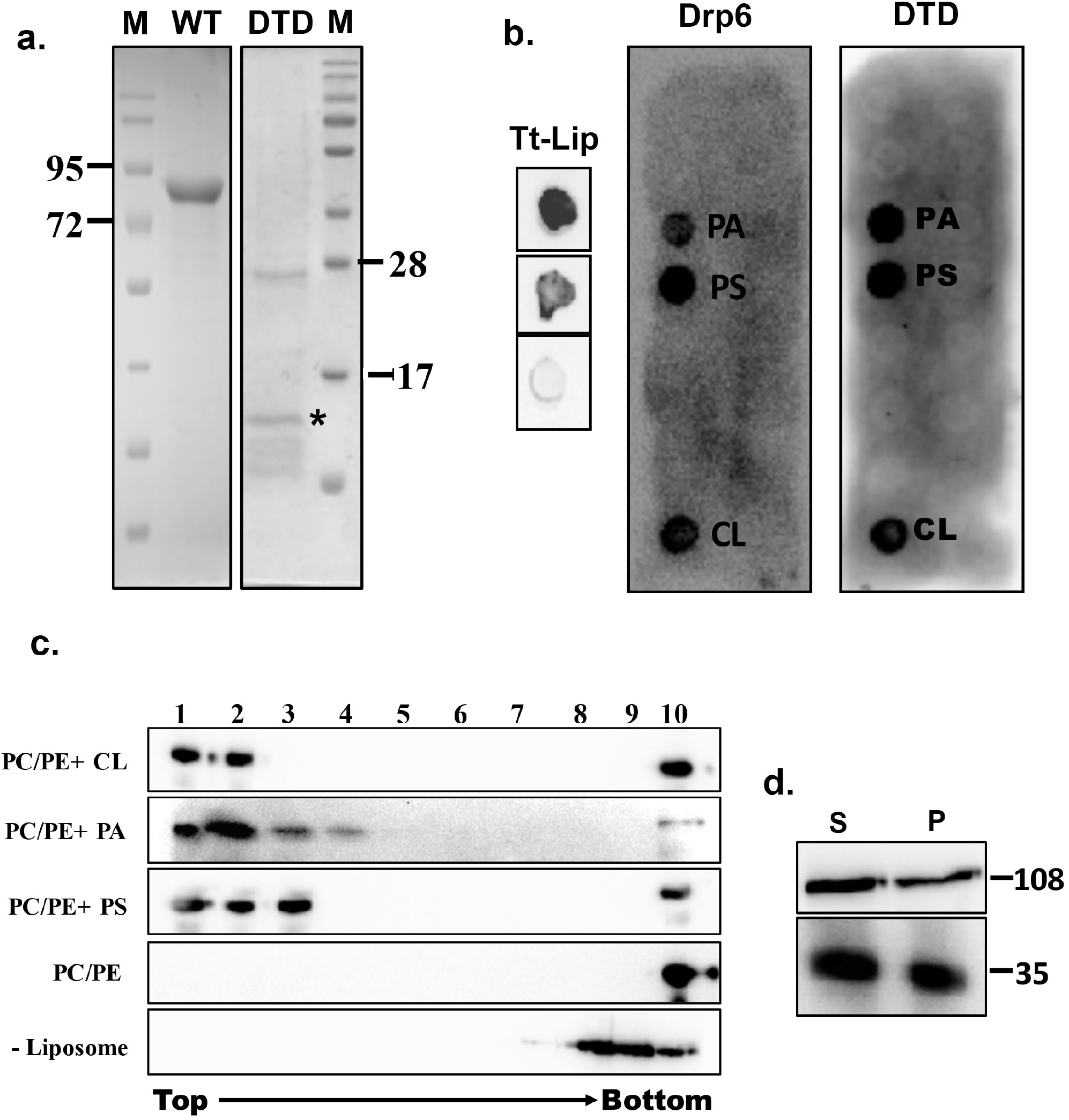
Identification of membrane-binding domain of Drp6. *a*. Coomassie stained SDS-PAGE gels showing purification of His-Drp6 (WT) and, His-DTD (DTD) expressed in *E. coli*. M is the molecular weight markers. Some of the markers are indicated on the sides. The purification of His-Drp6 DTD was partial and contained additional proteins from *E.coli* including one prominent band below 28kDa. The purified His-Drp6 DTD appearing below 17 kDa marker is indicated by an asterisk. *b*. Lipid overlay assay as detected by western blot analysis using anti-his antibody. (Tt-Lip); total *Tetrahymena* lipid spotted on nitrocellulose membrane and incubated with His-Drp6 in absence (top) or presence (middle) of GTP. The bottom spot is incubated with buffer without protein. Strip spotted with 15 different lipids and incubated either with His-Drp6 (Drp6) or with His-DTD (DTD). Both Drp6 and DTD interacted with Phosphatidic acid (PA), Phosphatidylserine (PS) and cardiolipin (CL) and are indicated. *c*. Floatation assay using liposomes containing 70% Phosphatidylcholine and 20% phosphatidylethanolamine additionally supplemented with 10% Cardiolipin (PC/PE+CL), 10% Phosphatidic acid (PC/PE+PA), 10% Phosphatidylserine (PC/PE+PS). While Liposomes in (PC/PE) contained 80% Phosphatidylcholine and 20% phosphatidylethanolamine, no liposome was added in (-Liposome). His-Drp6 was incubated either with different liposomes or without liposomes, overlaid with sucrose gradient and subjected to ultra-centrifugation. Fractions were collected from top and detected by western blot analysis using anti-his antibody. Drp6 appearing in the top four fractions indicate interaction with liposome. The experiments were repeated at least three times and representative results are shown here. *d*. Lysates of *Tetrahymena* cells expressing either GFP-Drp6 (Top panel) or GFP-Drp6 DTD (bottom panel) were fractionated into soluble (S) and membrane (P) fractions, and detected by western blot using anti-GFP antibody. Molecular weights of the proteins are indicated on the right.

We then looked at lipid binding in the physiologically-relevant context of a bilayer, using an *in vitro* binding assay to liposomes containing 10% PA, 10% PS, or 10% CL. All the liposomes also contained 70% PC and 20% PE. In sucrose density flotation gradients, the recombinant Drp6 co-migrated with all the three types of liposomes, appearing in the top (light) fractions (Fig. 2c). Drp6 was not found in the gradient top fractions in the absence of added liposomes (Fig. 2c). Similarly, Drp6 also failed to co-migrate with liposomes that contained only PC and PE (Fig. 2c). Taken together, our results indicate that Drp6 interacts with membranes *in vitro*, and that this depends on the presence of either PS, PA or CL. DTD by itself interacts with the same three phospholipids as holo-Drp6, consistent with the idea that DTD is the membrane-binding domain. To further test this idea, we performed sub-cellular fractionation of cells expressing *GFP-drp6-DTD* or *GFP-drp6.* As shown in Fig. 2d, Drp6 appeared in both soluble and membrane fractions. DTD also appeared in the membrane fraction, suggesting that it can bind to membranes *in vivo* (Fig. 2d). Taken together, these results lead us to conclude that DTD is a membrane-binding domain and requires phosphatidylserine, phosphatidic acid or cardiolipin for association with the membrane.

### A single point mutation (I553M) in the membrane-binding domain abrogates nuclear recruitment of Drp6

Dynamin binds to membrane lipids via interaction between its PH domain and the PIP2 head group. A hydrophobic patch in the PH domain of classical dynamin is important for membrane association/insertion (Ramachandran, Pucadyil et al. 2009). The sequence similarity between PH domain of human dynamin 1 and corresponding region of Drp6 is very low (Fig. 1b). However, the structural similarity is very high among all the dynamin family proteins whose structures are known. Therefore, we generated a three-dimensional model of Drp6 in order to identify the corresponding hydrophobic region in the DTD. The 3-D modelling of Drp6 shows that the structure of DTD is not related to PH domain, but that nonetheless has a hydrophobic patch (aa 553 - 555) (Fig. 3a). To test the importance of this patch, we substituted the first residue, I553, with M. We expressed this mutant allele as an N-terminal GFP-fusion (GFP-Drp6-I553M), and in parallel expressed a GFP-fusion of the wildtype protein (GFP-Drp6), and characterized their localization in *Tetrahymena* by confocal microscopy. While the latter localized mainly on the nuclear envelope with few cytoplasmic puncta, the mutated GFP-Drp6-I553M was not visible on the nuclear envelope but was instead exclusively present at cytoplasmic puncta (Fig. 3b). This difference suggests that the isoleucine at 553^rd^ position is important for nuclear localization of Drp6.

**Fig. 3.**
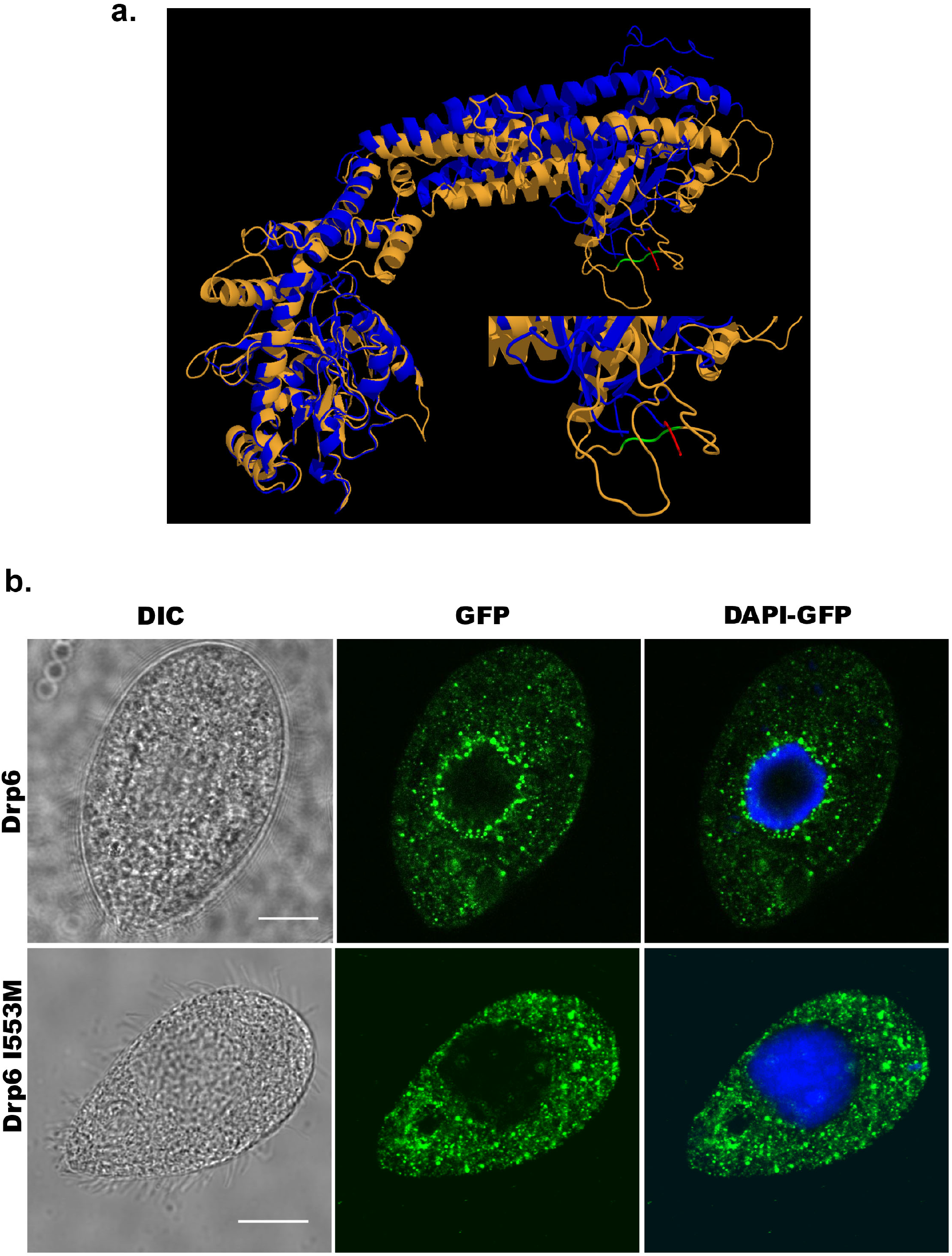
An isoleucine in the membrane binding domain is important for nuclear localization of Drp6. a. Three dimensional structure of Drp6. Homology model of Drp6 (brown) was generated by I-TASSER using Human Dynamin-1 as template (blue). The part containing the hydrophobic patch (red) in the PH domain of Human Dynamin-1 important for membrane insertion along with the putative hydrophobic patch (green) of Drp6 model are shown at the bottom right after enlarging the area. Although far apart in primary sequences, the regions containing hydrophobic patch in both the proteins come to the vicinity in 3-D structure. b. Localization of GFP-Drp6 (top) and GFP-Drp6 I553M (bottom). Confocal images of fixed *Tetrahymena* cells were obtained after DAPI staining. Mutation of isoleucine to methionine at 553^rd^ position leads to loss of nuclear localization. DAPI in blue shows the nucleus. Bar=10μm

### Mutation at I553 does not affect GTPase activity and self-assembly of Drp6

Dynamin and dynamin-related proteins require binding and hydrolysis of GTP for target membrane localization and membrane remodeling functions. Drp6 hydrolyses GTP *in vitro* (Kar, Dey et al. 2018). To understand if the mutation at I553 affects GTP binding and/or hydrolysis, we expressed and purified Drp6-I553M and Drp6 as N-terminal histidine tagged proteins (Fig. 4a) and compared their GTPase activities. The GTPase activity of Drp6-I553M (0.061± 0.002 nmol/μM/min) was not significantly different from that of Drp6 (0.056 ± 0.003 nmol/μM/min) (Fig. 4b). The GTPase activities of both wildtype and mutant proteins were also found to be similar when reactions were carried out for 0 to 20 min (Fig. 4b). We also assessed the Michaelis-Menten constant (K_m_) and maximum velocity (V_max_) for both these proteins (Fig. 4c). The K_m_ of Drp6-I553M (384 μM) was found to be slightly higher than that of Drp6 (180 μM) suggesting a marginal decrease in GTP binding affinity. The mutation did not affect V_max_ (0.089 nmol/uM/min for Drp6; 0.0893 nmol/uM/min for Drp6-I553M). Taken together, these results suggest that the defect in nuclear localization of Drp6-I553M is not due to defective GTPase activity.

**Fig. 4.**
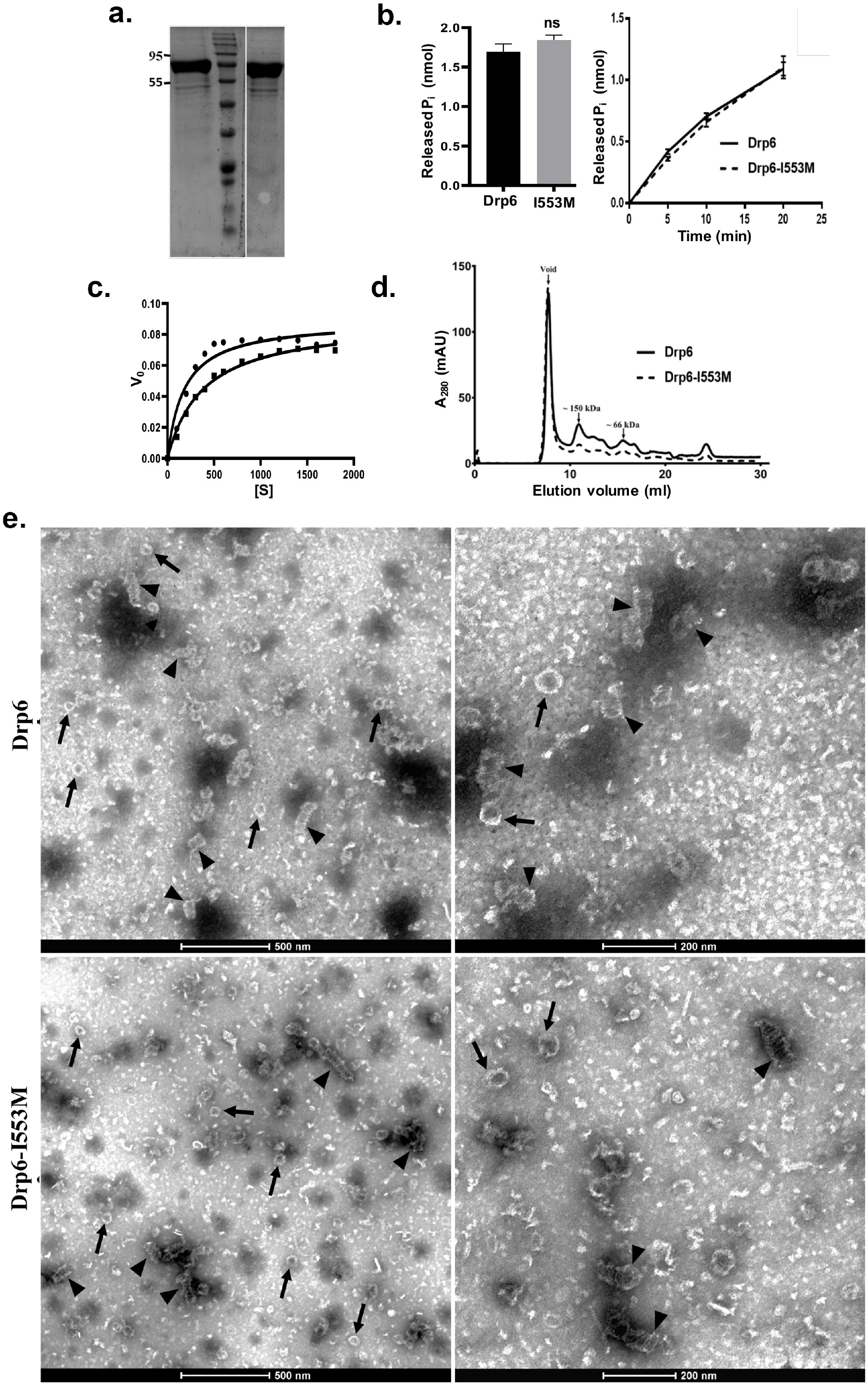
Mutation at I553 does not affect GTP hydrolysis activity and self-assembled structures. a. Coomassie stained SDS-PAGE gel showing purified His-Drp6 (lane 1) and His-Drp6 I553M (lane 3). Lane 2 is molecular weight marker. The positions of molecular weight are indicated on the left b. Graph showing GTP hydrolysis of Drp6 and Drp6-I553M as measured by phosphate release after 30 min of reaction (left). The graph on right shows reactions carried out for 0 to 20 min. c. Michaelis-Menten plot showing GTP hydrolysis by Drp6 (circle) and Drp6-I553M (square). V_0_= Rate of product formation in nmol P_i_/μM protein/min and [S]= GTP concentration in μM. d. Chromatograms depicting elution profiles of His-Drp6 and His-Drp6-I553M using superdex 200 size-exclusion column. The void volume and the positions of molecular weight markers are indicated by arrows. e. Electron micrographs of negatively stained His-Drp6 (Drp6) and His-Drp6-I553M (Drp6-I553M) at two different magnifications. Helical spirals and the ring structures are found in both wildtype and mutant proteins, and are indicated by arrow head and arrow respectively.

Another property of dynamin related proteins is their ability to assemble and disassemble at their target membranes. This involves self-assembly of helical spirals and ring structures, and is important for membrane association and membrane remodeling functions. The self-assembly of Drp6 and Drp6-I553M was evaluated by size exclusion chromatography. Drp6 eluted in the void volume as an oligomer containing at least 6 monomers, as previously observed (Kar, Dey et al. 2018) (Fig. 4d). Similarly, Drp6-I553M also formed higher order structures, as the majority of the protein eluted in the void volume (Fig. 4d). A small peak of material eluting near the 150 kDa marker might correspond to a dimer. The breadth of this smaller peak suggests it consists of a mixture of monomeric and oligomeric structures, with a dimer at the peak fraction. Since the mutant protein was able to form higher order oligomeric structure, we suggest that mutation does not inhibit its self-assembly. However, there might be a difference in the assembly products formed by the wildtype and mutant proteins. To examine this possibility, we compared the ultrastructure of the wildtype and mutant proteins by electron microscopy of negatively stained preparations of the purified recombinant proteins. The Drp6 appeared mostly as large helical spirals which are similar to structures found in other DRPs (Fig. 4e and Fig. S2). Ring-like structures were also present, as also found in other members of the family. Similarly, in Drp6-I553M samples we observed both helical spirals and ring-like structures (Fig. 4e, S2). These results suggest that the I553M mutation does not block in vitro oligomerization of Drp6. Taken together, our results suggest that defective nuclear localization of Drp6-I553M is not due to defects in GTPase activity or perturbation of self-assembly.

### Cardiolipin is important for nuclear recruitment of Drp6

Dynamin family proteins including Drp6 associate with their target membranes by interacting with specific lipids. To ask whether the I553M mutation might affect these interactions, we used *in vitro* membrane binding assays. Recombinant Drp6 and Drp6-I553M proteins were incubated separately with liposomes containing either 10% CL, PS, or PA. The association of the proteins with liposomes was then judged based on their co-flotation in sucrose density gradients. As can be seen in Fig. 5a, Drp6 co-floated with all the three liposomes and appeared on the top fractions, whereas Drp6-I553M co-floated with PS- or PA-containing liposomes but failed to co-float with CL-containing liposomes. Since the mutant retains the ability to associate with PS- and PA-containing liposomes, the overall membrane binding activity of Drp6 does not depend on I553. Instead, the mutation of I553 specifically inhibits interaction with CL. Based on these results, we infer that cardiolipin interacts with I553 in the membrane-binding domain, and that this interaction is important for membrane targeting in vitro.

**Fig. 5.**
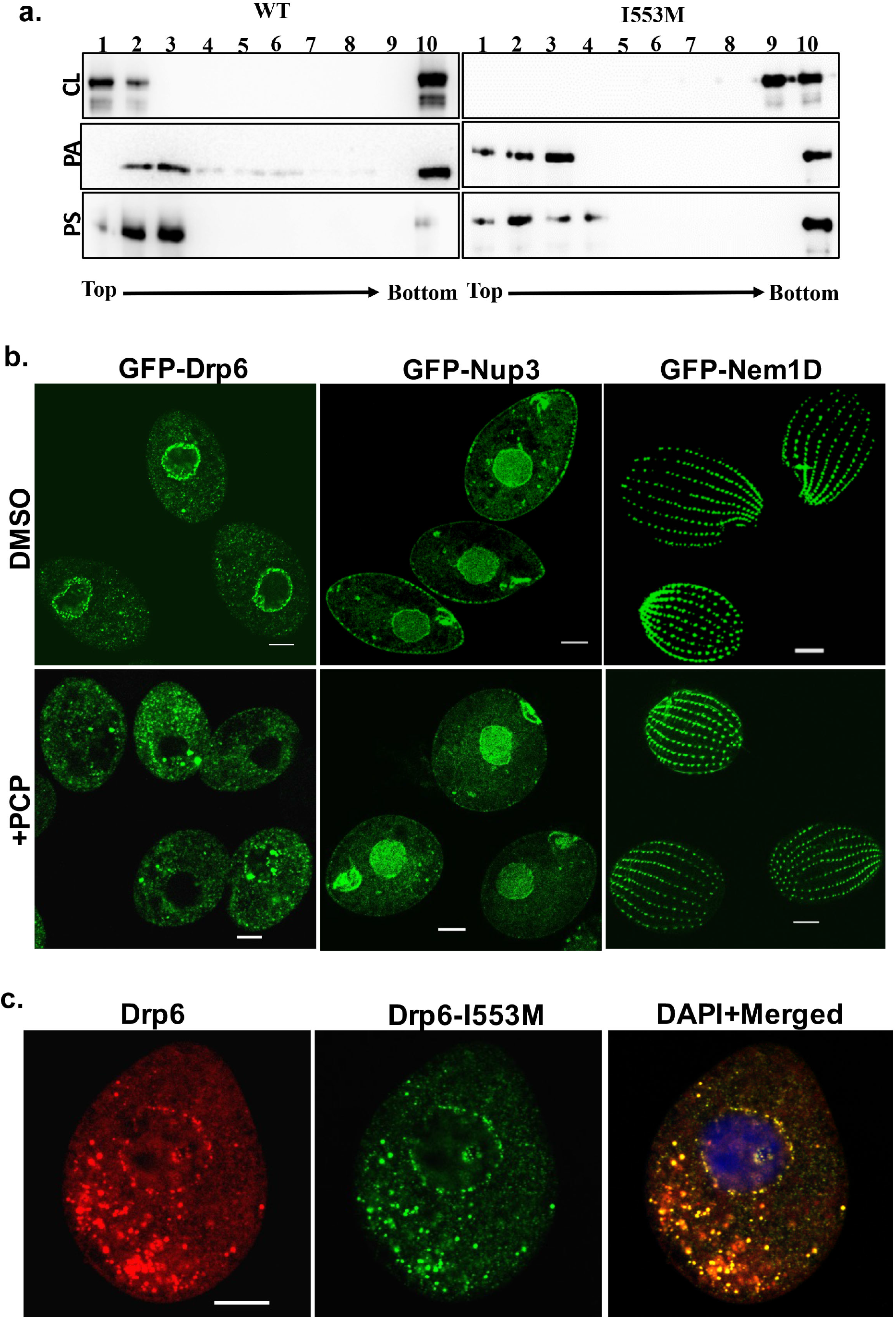
Interaction of cardiolipin with membrane binding domain recruits Drp6 to the nuclear membrane. a. Floatation assay was performed using liposomes with same composition and analysed by western blotting as mentioned in Fig. 2c. The assay was performed with wildtype Drp6 (WT) and Drp6-I553M (I553M). Liposomes supplemented with 10%Cardiolipin (CL), 10% Phosphatidic acid (PA), 10% Phosphatidylserine (PS) were used for the assay. Fractions collected from top to bottom are indicated. Experiments were repeated at least three times and representative results are shown here. Mutation at I553 lost interaction completely with the liposomes containing cardiolipin while retaining interactions with liposomes containing either phosphatidylserine or phosphatidic acid, suggesting isoleucine residue at 553^rd^ position is important for binding with cardiolipin in the bilayers. b. Confocal images of fixed *Tetrahymena* cells expressing GFP-Drp6 (left panel), GFP-Nup3 (middle panel), and GFP-Nem1D (right panel) either in presence (+PCP) or absence (DMSO) of PCP. Bar= 10μm c. Confocal images of fixed *Tetrahymena* cells co-expressing mCherry-Drp6 (left panel), and GFP-Drp6 I553M (middle panel). Merged image with DAPI stained nucleus is shown in right panel. Yellow colour in the merged image signifies presence of both Drp6 and Drp6-I553M in the same complex. Bar=10μm

We then asked whether the interaction between Drp6 and cardiolipin was important for nuclear targeting in vivo. Importantly, while the nuclear envelope of animals (Keenan, Berezney et al. 1970, Kleinig, Zentgraf et al. 1971, Sato, Fuji et al. 1972, Jarasch, Reilly et al. 1973) lacks cardiolipin, cardiolipin is present in the nuclear membrane of *Tetrahymena* (Nozawa, Fukushima et al 1973). To evaluate if cardiolipin is required for nuclear localization of Drp6, we depleted cardiolipin from *GFP-drp6* expressing cells using pentachlorophenol (PCP), a polychlorinated aromatic compound. PCP is a respiratory uncoupler and a potent inhibitor of cardiolipin synthesis (Ono and White 1971). Within 30 minutes of PCP treatment, GFP-Drp6 dissociated from nuclear envelopes in the majority of cells while remaining associated with cytoplasmic puncta (Fig. 5b, S3). Quantitative analysis showed that while GFP-Drp6 was localized at the nuclear envelope of all untreated cells, more than 80% of the cells completely lost nuclear localization of GFP-Drp6 with the remaining cells showing decreased nuclear localization upon PCP treatment. To check that the nuclear envelope itself remains intact under these conditions, we localized the nuclear pore protein GFP-Nup3. We found that GFP-Nup3 was clearly associated with nuclear envelopes before or after PCP treatment (Fig. 5b, S3). Moreover, PCP treatment does not disrupt membrane structure in general since the distribution of a cortical membrane-binding protein GFP-Nem1D (Shukla, Pillai et al. 2018) was also not affected by this treatment (Fig. 5b, S3). These results therefore suggest that the delocalization of GFP-Drp6 upon PCP treatment is due to loss of cardiolipin.

If the defect in localization of Drp6-I553M is due to a defect in CL-dependent targeting, one might expect that the defect would be suppressed in the presence of the wildtype protein, since the mutant protein would co-assemble with the correctly-targeted wildtype. To test this idea, we co-expressed *GFP-drp6-I553M* and *mCHERRY-drp6*. mCHERRY-Drp6 colocalized almost entirely with GFP-Drp6-I553M and was targeted to nuclear envelopes as well as cytoplasmic puncta, strongly suggesting that the mutant protein is able to co-assemble with the wildtype protein (Fig. 5c). This result also reinforces the earlier conclusion that the mutation does not affect the overall structure of the protein.

### Interaction of cardiolipin with Drp6 via membrane-binding domain is required for nuclear expansion

Previously, it has been shown that Drp6 is essential for MAC development in *Tetrahymena* (Rahaman, Elde et al. 2008). We have now established that interaction of Drp6 with cardiolipin is important for nuclear recruitment. Therefore, we hypothesized that inhibition of cardiolipin-Drp6 interaction would inhibit Drp6 function in MAC development. We took two independent approaches to perturb the interaction between cardiolipin and Drp6, and assessed the effect on MAC development. In the first approach, cardiolipin was depleted by treating cells at a stage prior to MAC development with PCP, and then measuring the efficiency of new MAC formation in conjugating cells (Fig. 6a). Quantitative analysis showed that while 71 ± 3.6% of the conjugants developed MACs in the control pairs, only 24 ± 1.7% developed MACs in the PCP-treated pairs (Fig. 6a). In the second approach, we perturbed cardiolipin-Drp6 interaction by treating conjugants with nonyl acridine orange (NAO). NAO interacts with cardiolipin with very high affinity and has been used in mammalian cells to block interactions between cardiolipin and mitochondrial proteins involved in electron transport (Maftah, Petit et al. 1990). Exposing conjugants to NAO significantly inhibited new MAC development (Fig. 6a).

**Fig. 6.**
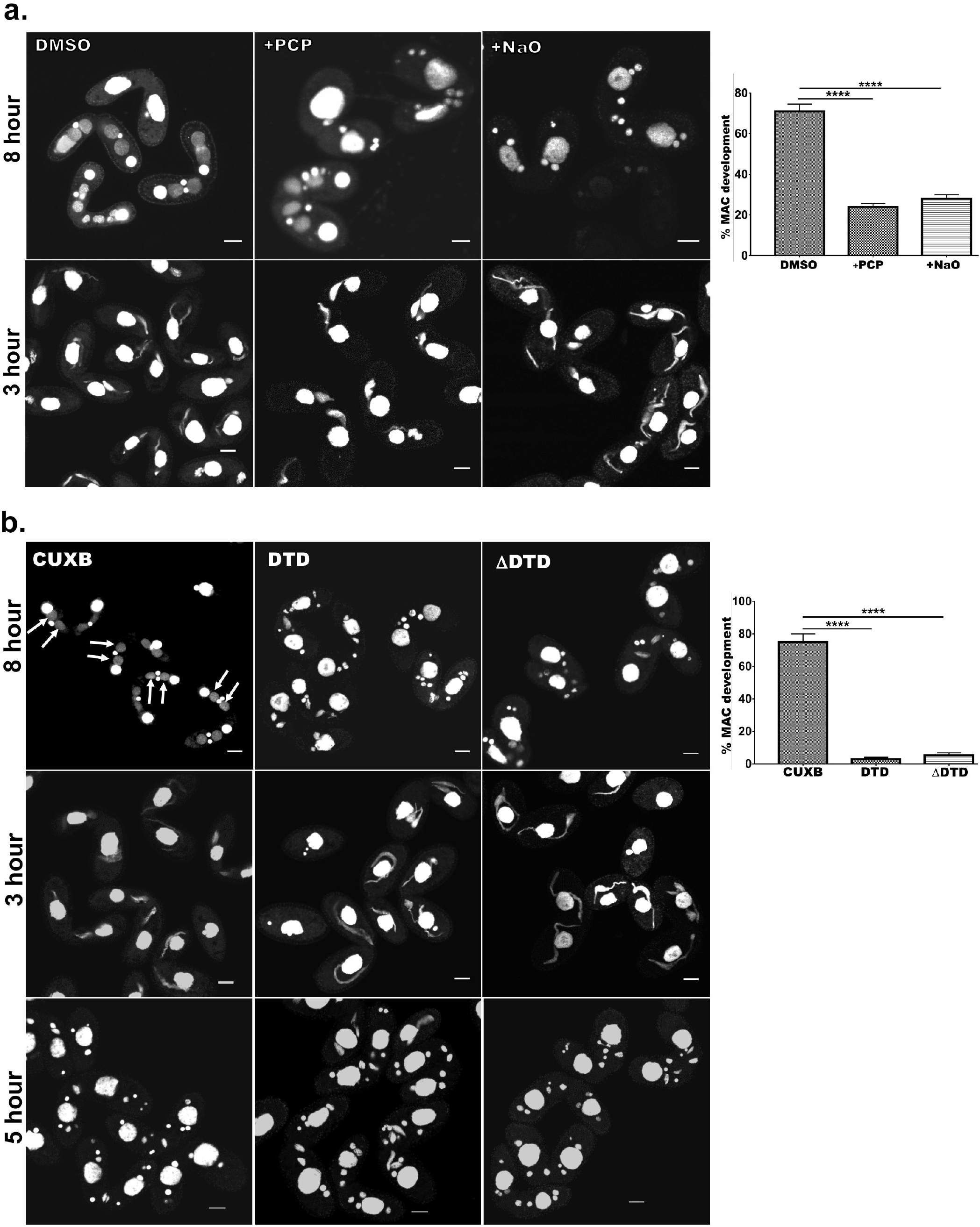
Cardiolipin and membrane binding domain regulate macronuclear expansion. a. Confocal images of fixed and DAPI stained conjugation pairs of *Tetrahymena* at 8 hours and 3 hours post conjugation. Two wildtype strains of *Tetrahymena* (Cu428 and B2086) were conjugated and treated either with pentachlorophenol (+PCP) or with nonyl acridine orange–D (+NaO) or with DMSO (DMSO). Top panel shows MAC development at 8 h and bottom panel shows MIC elongation at 3 h. Percent MAC development at 8 h is shown at the right. b. Confocal images of fixed DAPI stained *Tetrahymena* cells conjugated either between CU428 and B2086 (left panel) or between CU428 and GFP-Drp6 DTD (middle panel) expressing cells or between CU428 and GFP-Drp6 ΔDTD expressing cells (right panel). Top panel MAC development stage at 8 h, middle panel MIC elongation stage at 3 h and bottom panel meiotic stage at 5 h. The newly developed MAC is indicated by arrow. For quantitation, three independent experiments were performed and analyzed by unpaired T test (**** indicates p≤0.0001).

In conjugating *Tetrahymena*, MAC development is not the only phenomenon requiring nuclear expansion. At a prior stage, the germline micronuclei (MICs) show dramatic elongation (Cole, Cassidy-Hanley et al. 1997). We found that PCP treatment did not affect the frequency of elongation, i.e., the percentage of pairs showing elongated MICs, but did produce a decrease in the extent of MIC elongation (6A). This inhibition of elongation had no detectible consequences for the subsequent stage of MIC meiosis (Fig. 6a). Treatment with NAO had no measurable effect on MIC elongation or the subsequent meiosis (Fig. 6A). These results are consistent with the idea that the key requirement for cardiolipin is during MAC expansion.

We next reasoned that if interaction of Drp6 with cardiolipin is important for MAC expansion, then over-expression of the isolated Drp6 membrane-binding domain might competitively inhibit the interaction and block Drp6 function during MAC expansion. To test this possibility, we expressed *GFP-drp6-DTD* in *Tetrahymena* and then allowed the cells to conjugate with a Drp6 wildtype strain. We measured MAC development in these pairs, and in pairs from a parallel WT x WT cross, at 8 h during conjugation. In the control cross, more than 75% of pairs developed new MACs. In striking contrast, only 4-5% of pairs developed normal MACs in the pairs that included *GFP-drp6-DTD*-expressing cells (Fig. 6b). Therefore, MAC development was almost completely blocked when *GFP-drp6-DTD*-expressing cells comprised one of the conjugation partners (Fig. 6b).

In similar experiments, we also asked whether the expression of *GFP-drp6*Δ*DTD* might have a dominant-negative inhibitory effect on Drp6 function. Indeed, we found that in pairs where one cell over-expressed the ΔDTD construct, the pairs showed inhibition of MAC development that was similar to that induced by expression of the isolated DTD domain (Fig. 6b). Neither the expression of *GFP-drp6-DTD* nor *GFP-drp6*Δ*DTD* significantly reduced the fraction of pairs showing elongated MICs (Fig. 6b).

In conclusion, prior experiments with *DRP6* gene knockout pointed to a specific function in MAC development. Our current results from over-expression of mutant alleles are consistent with this idea. Taken together, our results support a model in which interaction of Drp6 with cardiolipin, for which a single amino acid acts as a key determinant, is critical for nuclear targeting and therefore for MAC expansion.

## DISCUSSION

Drp6 is a nuclear dynamin and is involved in nuclear remodeling (Rahaman, Elde et al. 2008). In the present study, we have identified a membrane-binding domain in Drp6 that directly interacts with lipids. Inhibition of cardiolipin synthesis blocks nuclear localization of Drp6, suggesting that it plays a critical role in Drp6 recruitment. Further evidence of cardiolipin interaction determining nuclear localization of Drp6 comes from the mutation of isoleucine at the 553^rd^ position. The mutant Drp6 protein loses its nuclear localization with concomitant loss of its interaction with cardiolipin without affecting other properties such as GTPase activity and self-assembly into rings/ helical spirals. The GFP-Drp6-I553M recruited to the nuclear envelope only when co-expressed with wildtype mCHERRY-Drp6, indicating that the mutant protein co-assembles with the wildtype (Fig. 3b). These results suggest that Drp6 molecules self-assemble on the nuclear envelope, and localizing the oligomer to the envelope does not require all subunits interact with cardiolipin. Taken together, these results suggest that the interaction between cardiolipin and the membrane-binding domain of Drp6 mediates the recruitment of Drp6 to the nuclear membrane.

We investigated the significance of cardiolipin in the nuclear remodeling function of Drp6. Inhibition of cardiolipin synthesis as well as perturbation of its interaction with Drp6 phenocopy the loss-of-function phenotype of Drp6, suggesting a role for cardiolipin in Drp6-mediated nuclear remodeling. Overexpression of Drp6-DTD or Drp6ΔDTD inhibits MAC expansion. Since DTD interacts with lipid including cardiolipin, the inhibition of MAC expansion is expected to be by competing with Drp6-cardiolipin interaction, concomitantly inhibiting Drp6 recruitment to the nuclear envelope. Inhibition of nuclear expansion by ΔDTD can be due to the inhibition of Drp6 localization on the nuclear envelope by forming a heterogenic complex as it lacks membrane-binding domain and does not associate with nuclear envelope. Based on these results, it can be concluded that cardiolipin acts as a molecular determinant in recruiting Drp6 on the nuclear envelope to perform nuclear remodeling function.

Drp6 is involved in nuclear expansion, which requires the incorporation of new lipids into the existing nuclear membrane, suggesting a membrane fusion function for Drp6. Consistent with this we recently observed that Drp6 is able to perform membrane fusion in vitro (our unpublished results). Membrane fission or fusion involves exchange of lipids between two juxtaposed bilayers, and optimum membrane fluidity is likely to be essential for the exchange of lipids between adjacent leaflets. Cardiolipin is known to facilitate the formation of apposed bilayers as well as to enhance membrane fluidity (Unsay, Cosentino et al. 2013). Dynamin proteins remodel membrane and bring bilayers to the vicinity during fission or fusion of membranes (Praefcke and McMahon 2004). Therefore, interaction of Drp6 with cardiolipin may enhance bilayer interaction and membrane fluidity. Taking together, it is reasonable to conclude that Drp6-cardiolipin interaction on the nuclear envelope facilitates nuclear expansion by enhancing membrane fusion and hence is essential for macronuclear expansion.

Drp6 interacts with three different lipids namely cardiolipin, phosphatidic acid and phosphatidylserine (present study). These interactions with multiple lipid might explain the localization of Drp6 in multiple sites (Fig. S4). The target specificity is often determined by the interaction of the membrane-binding domain with the specific lipids on the membrane. Although DRPs including Drp6 lack PH domain, they harbor a membrane-binding domain at the corresponding location (Fig. 1 and 2)(Ramachandran and Schmid 2018). The sequence diversity in this domain might explain the diverse functions of the family members on different target membranes. While PIP2 present in the plasma membrane associates with endocytic dynamin at the neck of vesicles, cardiolipin exclusively present in the mitochondria is recognized by the dynamins possessing mitochondrial remodeling function (Francy, Clinton et al. 2017, Kameoka, Adachi et al. 2018). Another lipid phosphatidylserine is abundantly present in mitochondria and is also recognized by mitochondrial dynamins (Yan, Qi et al. 2020). It is important to note that the nuclear envelope of *Tetrahymena* contains 3% cardiolipin (Nozawa, Fukushima et al 1973) and Drp6 (which is specifically present in the ciliate *Tetrahymena*) has evolved to interact with cardiolipin for specific recruitment to the nuclear envelope. In addition to nuclear envelope, Drp6 is also associated with ER vesicles (Rahaman, Elde et al. 2008). Localization of Drp6-I553M mutant on ER vesicles and its ability to interact with PS and PA on the membrane suggest that Drp6 localization on ER is dependent either on PA or PS or combination of both. Considering the abundance of PA on the ER membrane (Pillai, Shukla et al. 2017, Zegarlinska, Piascik et al. 2018) it could be argued that PA is involved in recruitment of Drp6 to ER. We also observed localization of Drp6 on plasma membrane (Fig. S4) and since PS is also present in the plasma membrane (Kay, Koivusalo et al. 2012), it is possible that Drp6 associates with plasma membrane via its interaction with PS. Although further experiments are required to find out the role of PS and PA in the recruitment of Drp6 in plasma membrane and ER, our results clearly show that lipid molecules play critical role in compartmentalizing the localization of Drp6 where CL shifts the dynamics from ER vesicles to nuclear envelope.

As mentioned earlier, I553 in the Drp6 membrane domain is critical for conferring specificity to Drp6 for recognizing cardiolipin on the nuclear envelope. The interaction of cardiolipin with I553 suggests the importance of hydrophobic patches for the target membrane specificity, since cardiolipin is known to interact strongly with hydrophobic residues (Planas-Iglesias, Dwarakanath et al. 2015). This is substantiated in the endocytic dynamin that uses a hydrophobic region including isoleucine at the 533^rd^ position in the PH domain for insertion into the target membrane (Ramachandran, Pucadyil et al. 2009). Our results on isoleucine mutation within the membrane domain also suggest the presence of a similar hydrophobic patch in Drp6 that is important for membrane interaction specifically via cardiolipin present on the nuclear envelope. However, hydrophobicity is not the sole determinant for the specific interaction with cardiolipin since methionine, which is also a strong hydrophobic residue, does not interact with cardiolipin when substituted for isoleucine (Fig. 3b). Therefore, it is conceivable that, in addition to hydrophobicity, the local conformation particularly the side chain of isoleucine plays a critical role in conferring the specificity for the recruitment to the target membrane via interaction with cardiolipin. Although residues important for target membrane selection have been identified in many dynamin proteins, they involve a stretch of positively charged amino acid for recognizing anionic head groups of several lipids including cardiolipin, phosphatidylserine and PIP2 (Salim, Bottomley et al. 1996, Achiriloaie, Barylko et al. 1999, Vallis, Wigge et al. 1999, Rujiviphat, Meglei et al. 2009, von der Malsburg, Abutbul-Ionita et al. 2011, Bustillo-Zabalbeitia, Montessuit et al. 2014, Smaczynska-de Rooij, Marklew et al. 2015, Wang, Guo et al. 2019). However, it is not known how different dynamin proteins distinguish different lipids solely based on ionic interaction. In the present study we demonstrate that a single isoleucine in the membrane-binding domain of Drp6 provides the specificity for cardiolipin. This isoleucine residue, however, does not influence Drp6 interaction with phosphatidylserine or phosphatidic acid, hence distinguishes among different negatively charged lipids. This is the first example of any dynamin protein in which a single amino acid site is shown to be important for conferring specificity to a lipid (cardiolipin), and thereby providing an additional target membrane (nuclear membrane) binding property to the protein. Perhaps this is also the first example in this family of proteins where a non-ionic interaction is shown to determine association with a specific target membrane. Although further experiments are needed to show the role of other two hydrophobic amino acid residues (M554 and I555) in the hydrophobic patch (aa 553-555 of Drp6) for the nuclear recruitment, our results provide the underlying mechanism of target membrane selection by a nuclear dynamin, and underscore the importance of cardiolipin interaction with a single amino acid residue in the Drp6 membrane binding domain in facilitating nuclear expansion.

## MATERIALS AND METHODS

### *Tetrahymena* strains and culture conditions

*Tetrahymena thermophila* CU428 and B2086 strains were obtained from *Tetrahymena* stock center, (Cornell university). Cells were cultured in SPP medium (2% proteose peptone (BD, USA), 0.2% glucose, 0.1% yeast extract and 0.003% ferric EDTA) at 30°C under shaking at 90rpm. For conjugation, mating type cells were grown in SPP media to a density of 3×10^5^ cells/ml, washed and resuspended in DMC media (0.17 mM sodium citrate, 0.1 mM NaH_2_PO_4_, 0.1 mM Na_2_HPO_4_, 0.65 mM CaCl_2_, 0.1 mM MgCl_2_) and incubated at 30°C, 90rpm for 16-18 hours. To initiate conjugation, starved cells of two different mating types were mixed and incubated at 30°C without shaking. All the reagents were purchased from Sigma Aldrich unless mentioned otherwise.

### Cloning and expression of transgenes in *Tetrahymena*

The Drp6ΔDTD lacking aa 517-600 was created by overlap PCR using a set of four oligonucleotides in a two-step PCR method. Drp6-I553M (mutation of isoleucine to methionine at 553^rd^ residue of Drp6) was generated by site-directed mutagenesis using Quick Change protocol (Stratagene). For expression in *Tetrahymena*, rDNA based vector pVGF or pIGF was used. While the PCR products of Drp6 (aa 1-710), Drp6ΔGED (aa 1-600) and Drp6-DTD (aa 517-600) were inserted between XhoI and ApaI restriction sites of pVGF, the Drp6-I553M and Drp6ΔDTD were introduced into pIGF using Gateway cloning strategy (Invitrogen) using the manufacturer’s protocol, and were expressed as N-Ter GFP tagged fusion proteins. All the constructs were confirmed by sequencing. Conjugating wildtype *Tetrahymena* cells were transformed with these constructs by electroporation, and the transformants were selected using 100μg/ml paromomycin sulphate. For the co-expression studies, *mCHERRY-Drp6* was generated by introducing mCHERRY sequences between PmeI and XhoI sites followed by Drp6 sequences between XhoI and ApaI sites of NCVB vector. Co-transformants were generated by biolistic transformation of linearized mCHERRY-Drp6 nucleotide sequences into the cells expressing *GFP-drp6-I553M*, and were selected in presence of 60 μg/ml blasticidine and 120 μg/ml paromomycin sulfate supplemented with 1 μg/ml cadmium chloride.

Cells were grown to a log phase (2.5 to 3.5×10^5^ cells/ml) and expression was induced by adding cadmium chloride at concentration of 1μg/ml for 4 hours. Cells were harvested at 1100 g, fixed with 4% paraformaldehyde for 20 minutes at RT, washed and resuspended in 10mM HEPES, pH 7.5 before imaging.

### Cloning, expression and purification of recombinant proteins in *E.coli*

For expression in *E. coli*, the amplified PCR products of Drp6, Drp6-I553M and Drp6-DTD were cloned into pRSETB using BamHI and EcoRI sites. The resulting constructs were transformed into chemically competent *E.coli* C41(DE3) cells and transformants were inoculated into LB broth supplemented with 100μg/ml ampicillin and grown at 37°C till the OD600 reached 0.4. The cultures were then shifted to 18°C and expression was induced after 1 hour by adding 0.5mM IPTG (Sigma) and kept for16 hours at the same temperature before harvesting the cells. The harvested cells were resuspended in ice cold Buffer A (25mM HEPES pH 7.5, 300mMNaCl, 2mMMgCl2, 2mM β-mercaptoethanol and 10% glycerol) supplemented with EDTA-free protease inhibitor cocktail (Roche) and 100mM phenyl methyl sulfonyl fluoride, lysed by sonication and the lysates were centrifuged at 52000 g for 45 min at 4°C.The supernatants were incubated with Ni-NTA agarose resin (Qiagen, Germany) for 2 hours before washing with 100 bed volume buffer A supplemented with 50mM imidazole. The bound proteins were eluted with 250mM imidazole in buffer A. The purified proteins were checked by Coomassie-stained SDS-PAGE gels and the purity was assessed by Image J analysis (NIH). The fractions containing the purified proteins were pooled, dialyzed with buffer A and concentrated using Amicon ultra-15 filters (Millipore). Protein concentration was estimated by Bradford assay (Bio-Rad Laboratories, USA).

### Western blotting

Samples were subjected to SDS-PAGE gel and the proteins were transferred to PVDF membrane. Membrane was blocked with 2% BSA in TBST (50 mM Tris-Cl, 150 mMNaCl, pH 8.0 and 0.05% Tween 20) for 1 hour. The blot was then incubated with HRP-conjugated anti-His monoclonal antibody (1:5000) and detected with supersignalfemto substrate (Thermoscientific, USA) using ChemiDoc imaging system (Bio-Rad laboratories, USA).

### Fractionation of membrane protein and soluble protein

*Tetrahymena* cells expressing GFP-Drp6 or GFP-DTD were lysed in 500 μl of ice cold lysis buffer containing 25mM Tris-Cl pH 7.5, 300mM NaCl, 10%glycerol supplemented with E-64, pepstatin, aprotinin, PMSF and protease inhibitor cocktail (Roche) by passing through a ball bearing homogeniser with a nominal clearance of 0.0007 in. The resulting lysates were centrifuged at 16000 g for 15 min at 4°C, the supernatant was collected as soluble protein fraction, and the pellet containing the membrane fraction was resuspended in 500 μl lysis buffer. The proteins in both soluble fraction and membrane fraction were separated in 12% SDS-PAGE gel and analysed by western blotting using anti-GFP polyclonal antibody (1:4000; Sigma-Aldrich).

### Lipid overlay assay

Total *Tetrahymena* lipid was extracted from growing *Tetrahymena* cells (5×10^5^ cells/ml) by method Bligh and Dyer (1959). Drops of 5 ul in chloroform were spotted on the nitrocellulose membrane and incubated with His-Drp6 (90 ug/ml) in GTPase assay buffer in presence or absence of 1 mM GTP for 1 hour. In control experiments, BSA (2) was used in place of His-Drp6. The assay using membrane lipid strips (P-6002, Echelon Biosciences, USA) spotted with 100 pmol of 15 different lipids were used according to the manufacturer’s instruction. The binding of proteins was detected by western blot analysis using anti-His monoclonal antibody.

### Floatation assay

Lipids (Avanti Polar) were dissolved in analytical grade chloroform, liposomes were prepared using 2.5 mg total lipid in 1ml chloroform. The liposomes contained 70% PC and 20% PE along with either 10% CL or 10% PA or 10% PS. A thin dry film was obtained by drying the solution in a round bottom flask and solvent was completely removed in a lyophilizer. Liposomes were made by rehydrating the film in buffer A (25mM HEPES pH 7.5, 2mM MgCl2, 150mM NaCl) pre-warmed at 37°C. The resuspended solution was extruded 17-21 times through extruder (Avanti Polar) using filter with100nm pore and the size distribution was measured by DLS in Malvern Zetasizer Nano. For floatation assay, 1μM protein was incubated with 0.5mg liposomes in buffer A supplemented with 1mM GTP for 1 hour at RT. Sucrose was added to the reaction mixture (final sucrose concentration 40%), placed at the bottom of a 13 ml ultra-centrifugation tube and overlaid with 2 ml each of 35%, 30%, 25%, 20%, 15% and 0% sucrose solutions in the same buffer. The gradient was subjected to ultra-centrifugation in Beckman Coulter ultra-centrifuge at 35,000 RPM for 15 hours at 4°C using SW41 rotor. Fractions (500μl) were collected from top and detected by western blotting using anti-His HRP conjugated monoclonal antibody (1:5000).

### Measurement of GTP hydrolysis activity

The GTP hydrolysis activity of recombinant Drp6 and Drp6 I553M was measured in a colorimetric assay using Malachite Green-based phosphate assay reagent (BIOMOL Green, Enzo Life Sciences). The GTPase assay (20 ul in 25mM HEPES pH7.5, 15mM KCl, 2mM MgCl_2_) was performed in presence of 1mM GTP (Sigma) using 1μM protein for 0-30 min at 37°C. The reaction was stopped by adding 5μl of 0.5 mM EDTA and absorbance was measured at 620 nm. For measuring Km and Kcat, reactions were performed for 10 min at 37°C in triplicate using varying concentrations of GTP (50 μM to 2000 μM). The values obtained from three independent experiments were plotted and analyzed using GraphPad Prism7 software.

### Size exclusion chromatography

Size exclusion chromatography was performed on the Superdex 200 GL 10/300 column (GE Life Sciences) using Akta Explorer FPLC system (GE healthcare) which was calibrated with standard molecular weight markers (Sigma). Five hundred microliters of protein (0.5mg/ml) in buffer A was loaded onto the pre-equilibrated column and was run at 0.5ml/min. The chromatogram was recorded by taking absorbance at 280 nm.

### Electron microscopy

Purified recombinant Drp6 or Drp6 I553M (1μM) was incubated with 0.5mM GTPγS in 25 mM HEPES pH 7.5, 150 mM NaCl and 2 mM MgCl_2_ for 20 min at room temperature and was adsorbed for 5 min onto a 200 mesh carbon coated Copper grid (Ted Pella, Inc.). The grid was stained with a drop of 2% freshly prepared uranyl acetate (MP Biomedicals, USA) for 2 min and dried at room temperature for 10 min. The electron micrographs were collected on a FEI Tecnai G2 120KV electron microscope.

### Confocal microscopy

*Tetrahymena* cells were fixed with 4% paraformaldehyde (PFA) in 50mM HEPES pH 7.5 for 20 min at RT and were collected in 10 mM HEPES pH 7.5 after centrifugation at 1100 X g. The fixed cells were stained with DAPI (0.25 ug/ml) (Invitrogen) and washed with 10 mM HEPES pH 7.5 before imaging. The images were collected in a Zeiss LSM780 or Leica DMI8 confocal microscope.

### Pentachlorophenol (PCP) and Nonyl acridine orange (NAO) treatment

The growing *Tetrahymena* cells either expressing *GFP-Drp6* or *GFP-Nup3* or *GFP-Nem1D* were treated with 10 μM PCP in DMSO for 30 minutes before fixing with 4% paraformaldehyde. The conjugation pairs were either treated with 30 μM PCP or 0.5 μM NAO (Invitrogen) after 2.5 h, 4.5 h or 7.5 h post-mixing, and were fixed after 30 minutes of addition. The cells were stained with DAPI (0.25 ug/ml) before imaging.

## Author Contributions

A.R. designed the experiments. U.P.K. and H.D performed all experiments. U.P.K., H.D. and A.R. analyzed the results. U.P.K., H.D. and A.R. wrote the manuscript.

## Acknowledgements

We thank Prof. Aaron Turkewitz from the University of Chicago, Dr Kausik Chakraborty from IGIB, Prof. Jacek Gaertig from University of Georgia and Dr Utpal Nath from Indian Institute of Science for critical evaluation and useful comments on the manuscript. The work was partly funded by DBT grant (BT/PR14643/BRB/10/862/2010) to AR. The funders have no role in study design and interpretation, or the decision to submit the work for publication.

## Abbreviations used are

DRP: dynamin related proteins
MIC: micronucleus
MAC: amcronucleus
DTD: Drp targeting determinant
PA: phosphatidic acid
PS: phosphatidylserine
CL: cardiolipin
ER: endoplasmic reticulum.

