## Supplemental Figures for "Cardiolipin targets a dynamin related protein to the nuclear membrane"

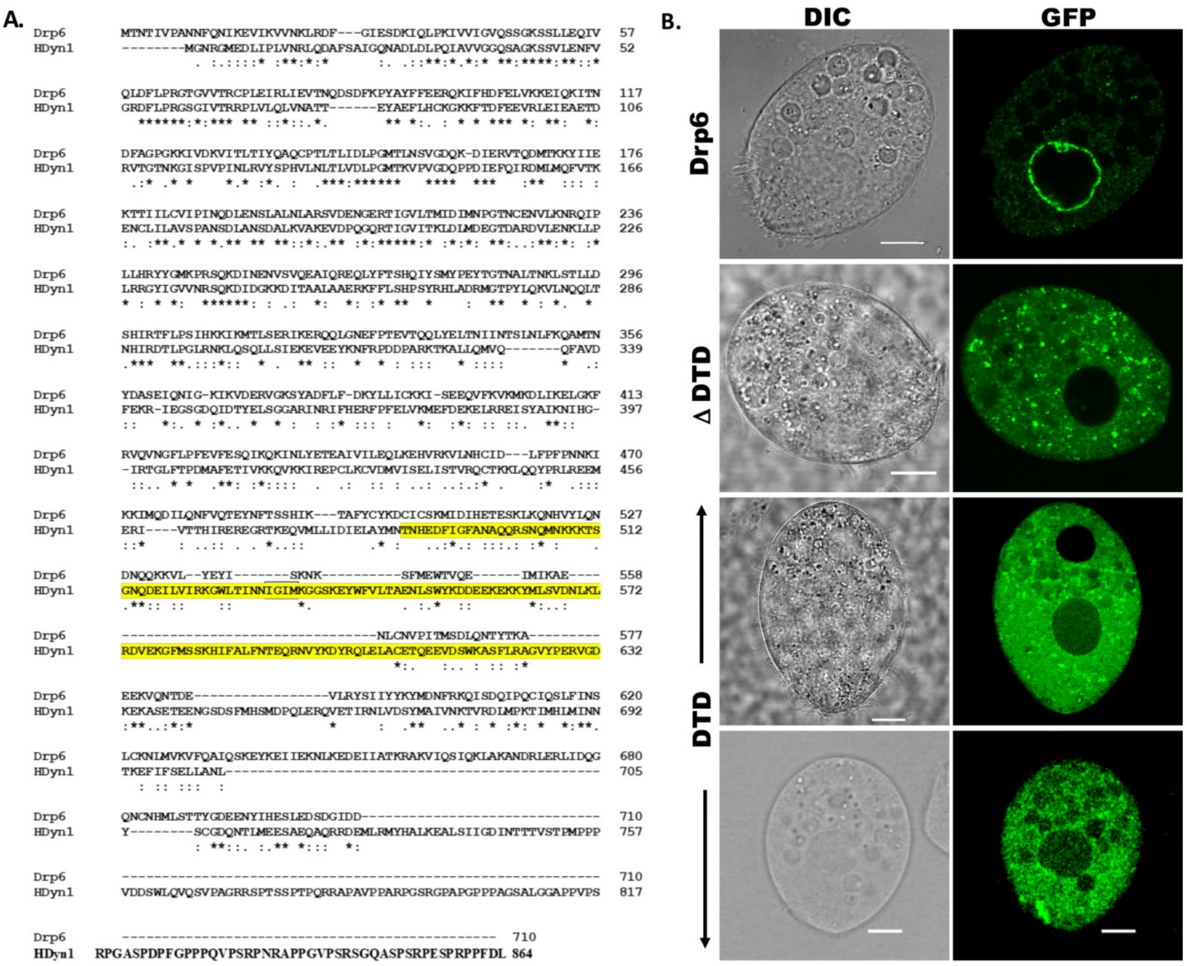


**Figure S1:** (A) Alignment of amino acid sequences of *Tetrahymena* Drp6 (Drp6) and human dynamin 1 (HDyn1). The PH domain of HDyn1 is highlighted in yellow.
(B) Confocal images of live *Tetrahymena* cells expressing GFP-Drp6 (Drp6), GFP-Drp6 ΔDTD (ΔDTD) and GFP-DTD (DTD). The similar distribution of expressed proteins in fixed cells (Fig. 1) shows that the fixation did not affect overall distribution. Bar = 10μm

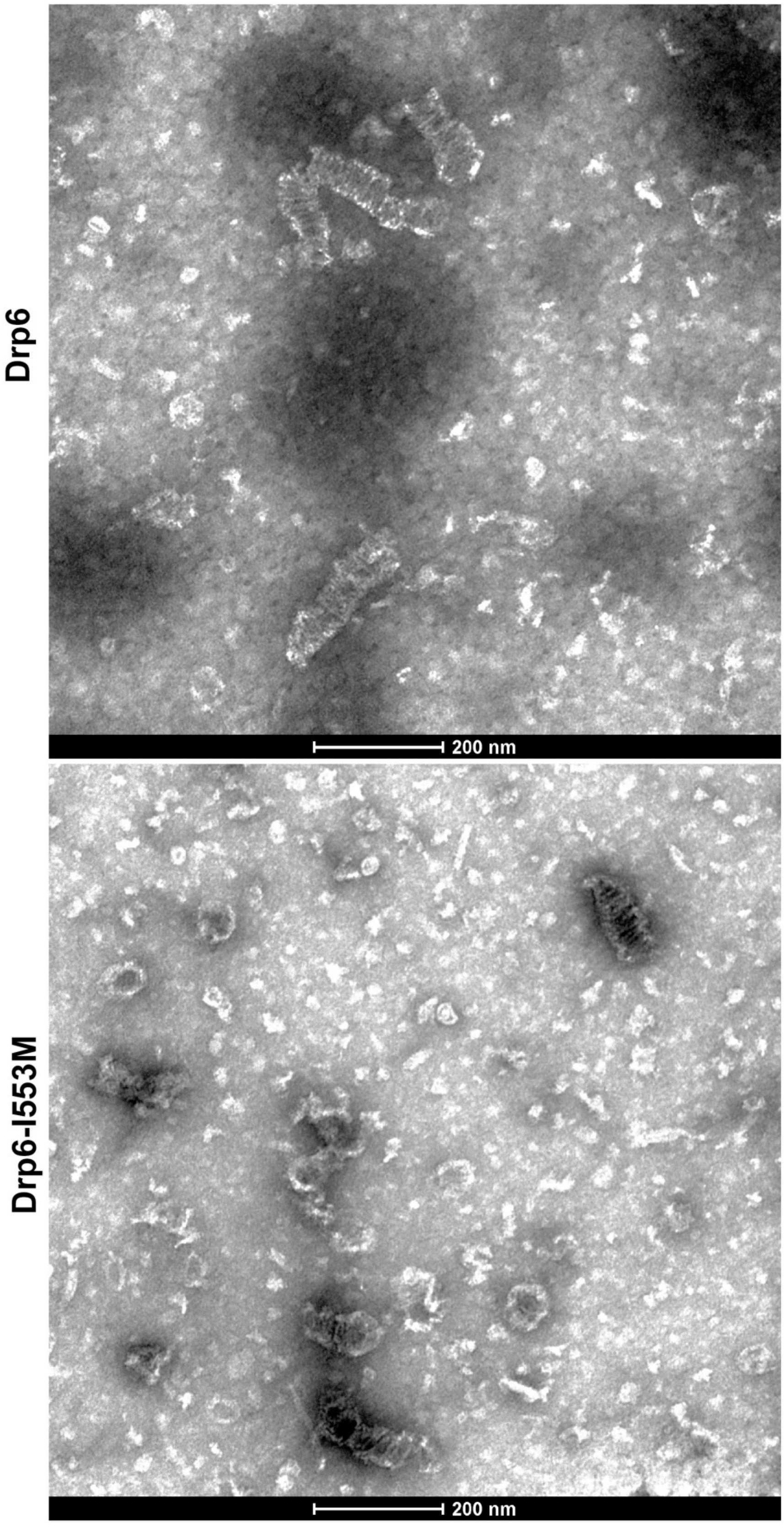


**Figure S2:** Electron micrographs of His-Drp6 (Top) and His-Drp6-I553M (Bottom) after negative staining.

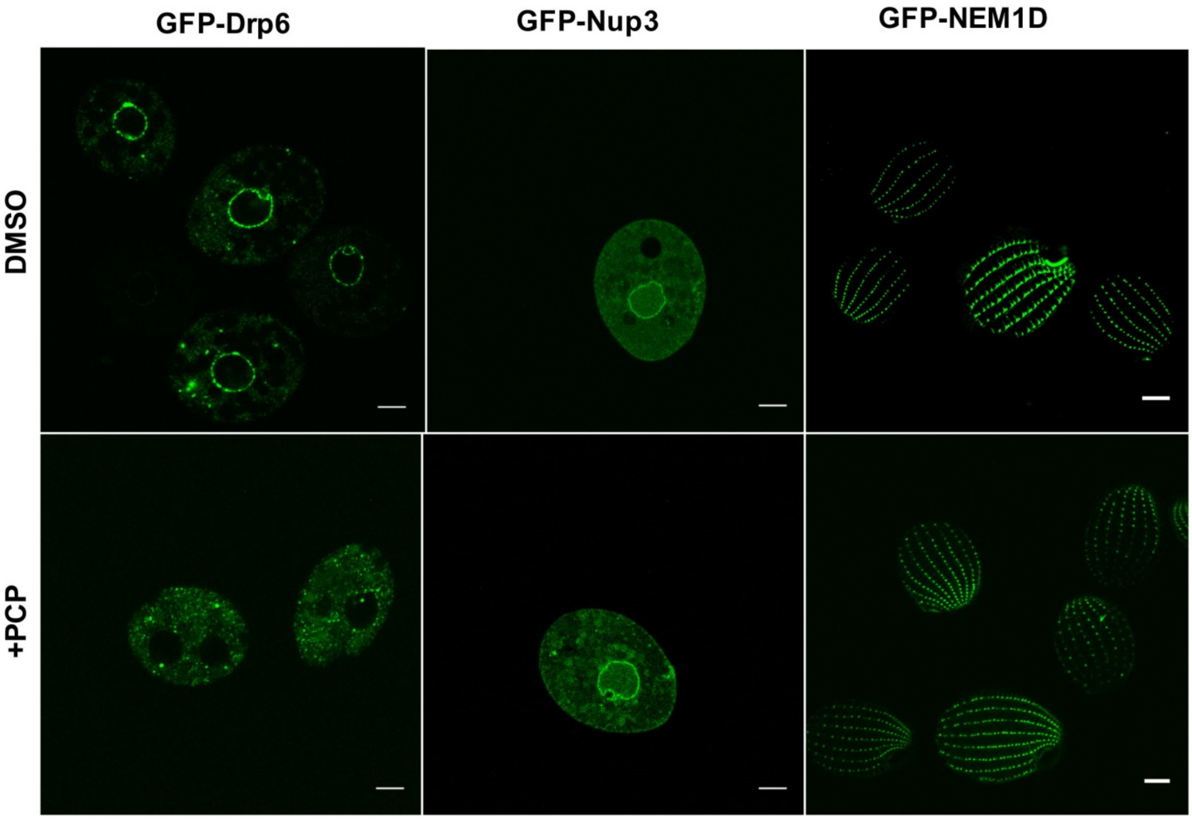


**Figure S3:** Confocal images of live *Tetrahymena* cells expressing GFP -Drp6 (left panel), GFP-Nup3 (middle panel), and GFP-Nem1D (right panel) either in presence (+PCP) or absence (DMSO) of PCP. Bar= 10μm

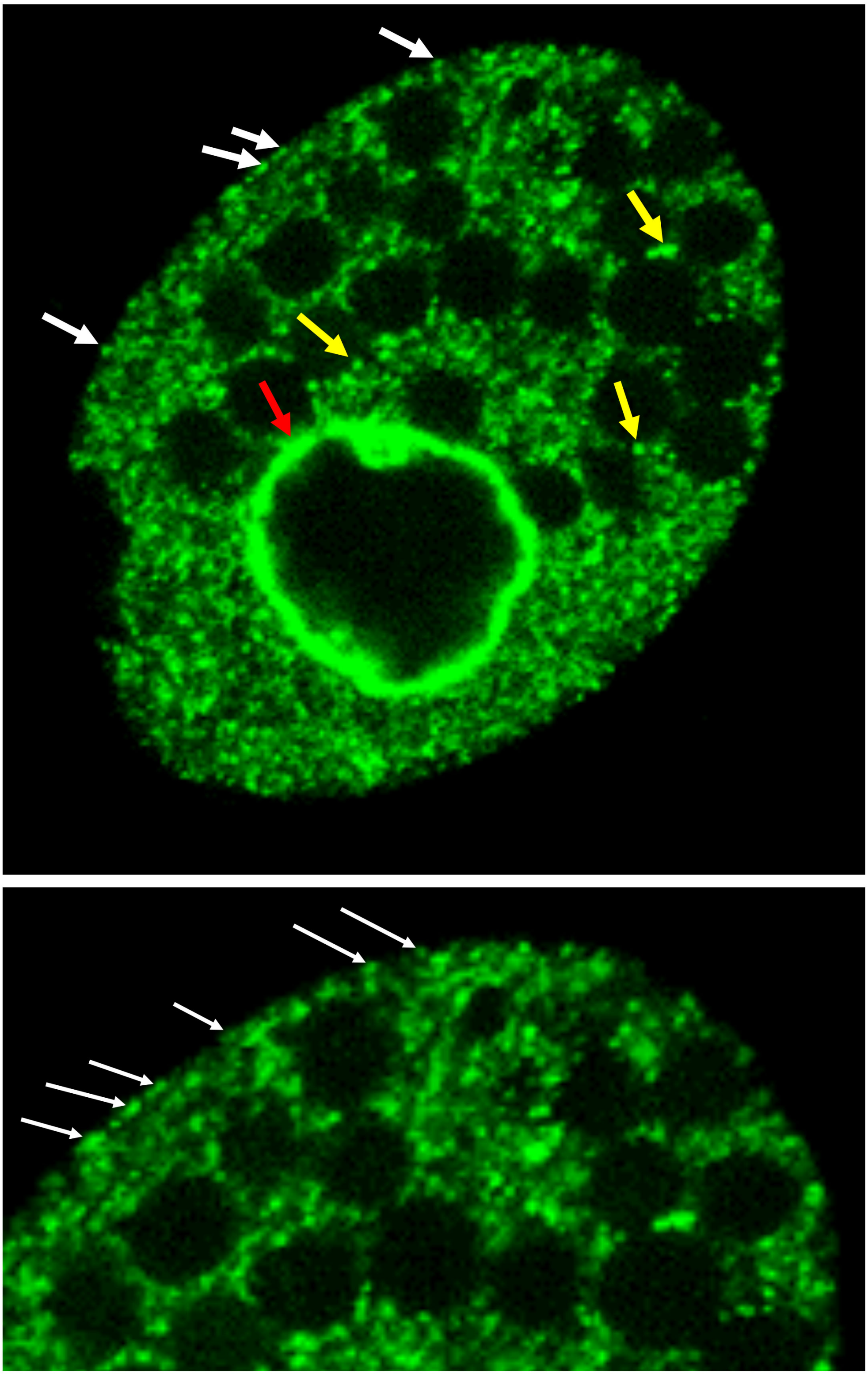

Figure S4: Dynamin related protein 6 (Drp6) localizes to multiple sites: Top: Confocal image of a live *Tetrahymena* cell expressing Drp6 as GFP fusion protein. The localization of Drp6 on nuclear envelope (red arrow), on ER vesicles (yellow arrow) and on plasma membrane (white arrow) are shown. Bottom: Part of the image is enlarged to visualize the localization on the plasma membrane more distinctly.
